# Super-resolution imaging of proteins inside live mammalian cells with mLIVE-PAINT

**DOI:** 10.1101/2024.07.22.604574

**Authors:** Haresh Bhaskar, Zoe Gidden, Gurvir Virdi, Dirk-Jan Kleinjan, Susan J. Rosser, Sonia Gandhi, Lynne Regan, Mathew H. Horrocks

## Abstract

Super-resolution microscopy has revolutionized biological imaging, enabling the visualization of structures at the nanometer length scale. Its application in live cells, however, has remained challenging. To address this, we adapted LIVE-PAINT, an approach we established in yeast, for application in live mammalian cells. Using the 101A/101B coiled-coil peptide pair as a peptide-based targeting system, we successfully demonstrate the super-resolution imaging of two distinct proteins in mammalian cells, one localized in the nucleus, and the second in the cytoplasm. This study highlights the versatility of LIVE-PAINT, suggesting its potential for live-cell super-resolution imaging across a range of protein targets in mammalian cells. We name the mammalian cell version of our original method mLIVE-PAINT.

## Introduction

In conventional microscopy, the overlapping fluorescence from molecules separated by distances shorter than the diffraction limit of light, ∼250 nm, prevents them from being distinguished (Abbe, 1873, Lelek et al., 2021). To overcome this limitation, super-resolution (SR) microscopy has been developed to study proteins at the nanometer length scale in various biological systems, achieving resolutions as high as 5 nm (Loschberger et al., 2014). One branch of SR microscopy, single-molecule localization microscopy (SMLM), achieves this by separating and locating the fluorescence from individual emitters over time as they stochastically blink (Lelek et al., 2021) (for a comprehensive review, see Horrocks et al., 2014).

In recent years, DNA-PAINT (Point Accumulation for Imaging in Nanoscale Topography) has emerged as a popular SMLM method for imaging protein targets in fixed biological samples. This approach involves tagging proteins of interest (POIs) with short DNA oligonucleotide “docker” strands. An image is constructed as fluorescently-tagged complementary “imager” oligonucleotides transiently hybridize with the docker strands and are localized with nanometer precision. DNA-PAINT, however, usually requires the use of antibodies or nanobodies (Schnitzbauer et al., 2017, Chung et al., 2022) as well as internalization of DNA imager strands, and so cannot be used to image internal structures in live cells.

Inspired by DNA PAINT, we developed an imaging strategy using peptide-peptide interactions that can be used in living cells. We demonstrated its efficacy in live yeast (Oi et al., 2020), Gidden et al., 2023). Analogous to DNA-PAINT, we demonstrated that a protein-of-interest (POI) tagged with one partner of a coiled-coil (CC) peptide pair can be localized as the partner CC peptide fused to a fluorescent protein (FP) transiently binds to it. We also recently imaged F-actin inside live mammalian cells using direct-LIVE-PAINT, which uses a peptide that transiently binds directly to the POI (Bhaskar et al., 2023).

In this study, we present mammalian LIVE-PAINT (mLIVE-PAINT) using the 101A/101B (Chen et al., 2015) CC peptide pair to image nuclear (H2B) and cytoplasmic (TOM20, mitochondria) targets in live SH-SY5Y cells. By generating stable cell lines expressing the FP-tagged CC imaging construct, mNeonGreen-101A, and transiently transfecting in our POI-101B, we imaged these targets using both spinning disk confocal microscopy (diffraction limited, DL) and total internal reflection fluorescence (TIRF) microscopy (SR). As a proof-of-principle, we demonstrate that mammalian LIVE-PAINT can be used to image both static and dynamic targets within living cells, offering versatile and minimally perturbative SR detection.

## Results

### 101A-101B peptide pair can be used to image proteins in mammalian cells

To use peptide pairs for imaging proteins in cells, they must bind to each other specifically to avoid mislocalization, and must also have a reasonably tight binding affinity (Kd in the nanomolar range). The CC pair 101A/101B is a 42 AA-long peptide pair published as part of a synthetic toolkit for barcoding yeast strains in high-throughput screening studies (Chen et al., 2015), and satisfies these two criteria. The LIVE-PAINT system using this pair consists of two components: (i) the 101A peptide fused to the C-terminus of the FP, mNeonGreen (mNG), stably expressed under a doxycycline inducible promoter in SH-SY5Y cells and (ii) its partner peptide 101B, fused to the C-terminus of the POI, transiently transfected into this stable line (Figure 1A,B).

**Figure 1.**
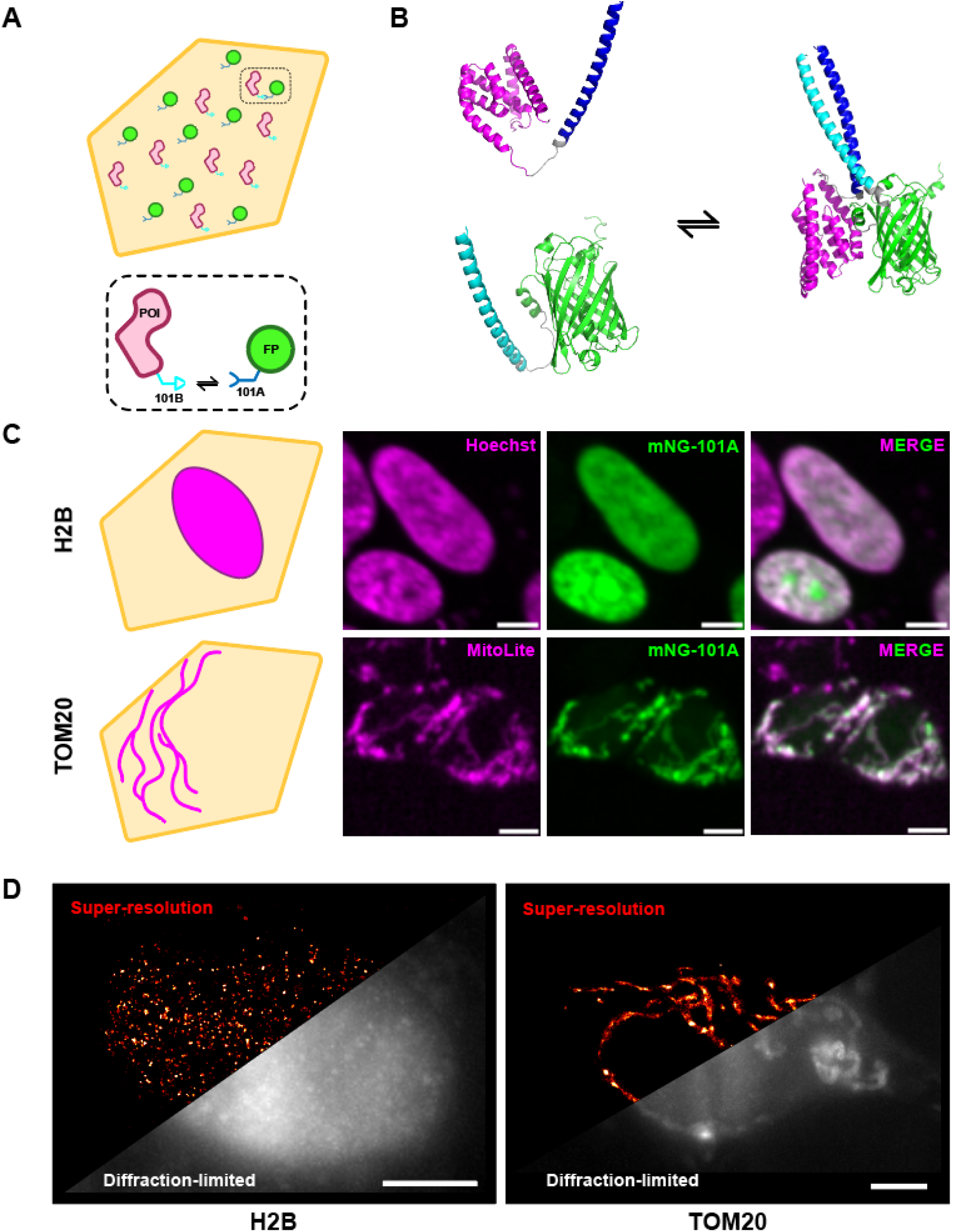
**(a)** Schematic representation of a mammalian cell expressing the components necessary for LIVE-PAINT imaging. Protein of Interest (POI) is shown in pink, fused to the 101B peptide (light blue) and the imaging component (mNeonGreen (mNG) fused to the 101A peptide) is shown in green and dark blue. **(b)** AlphaFold 3 structure prediction of mNeonGreen-101A and TOM20-101B and modelling of interaction via 101A/B peptide pair. mNeonGreen (green), 101A (cyan), TOM20 (magenta), 101B (blue) and linkers shown in gray. **(c)** Schematic representations of expected cellular locations and structures of nuclear protein H2B and mitochondrial protein TOM20 shown in the first column. Spinning disk confocal images of respective targets imaged with live cell organelle specific dyes (mitochondria - MitoLite Red FX600; nucleus - Hoechst) and the LIVE-PAINT system in SH-SY5Y cells transiently transfected with either TOM20-101B or H2B-101B shown in subsequent panels and overlayed. 5 μm scale bar. **(d)** Super-resolution (SR) reconstructions (colored ‘red hot’) of SH-SY5Y cells transfected with either TOM20-101B or H2B-101B and imaged in TIRF. Localisations were consolidated from 100 s of imaging (2000 frames, 50 ms exposure) and diffraction limited section of the field-of-view shown as a maximum intensity z-projection over the same imaging period. Precision threshold <30 nm. 5 μm scale bar.

We first assessed whether the 101A/101B peptide pair could effectively target and localize the imaging construct (mNG-101A) specifically to the POI (POI-101B). To test this, we transfected the mNG-101A-expressing stable cell line with TOM20-101B (a mitochondrial protein) or H2B-101B (a nuclear protein) and performed live imaging using spinning disk confocal microscopy (Figure 1C). The mNG signal showed specific localization to the mitochondria and nucleus under their respective transfection conditions, with high colocalization to their respective organellar stains (Pearson’s correlation H2B - 0.85, TOM20 - 0.89).

Next, we imaged these samples using TIRF microscopy to detect single binding events between the imager and POI with a higher signal-to-background ratio. Temporally separated blinking events were detected for both targets at an average localization rate of 250 localizations per second and used to reconstruct a SR image (Figure 1D). These results suggest that the 101A/101B peptide pair has suitable binding kinetics for single-molecule localization detection of both cytoplasmic and nuclear protein targets.

### Increased spatial resolution allows detection of ‘clutch’-like organisation of H2B

Increased spatial resolution can reveal structural organization at the nanometer scale that is undetectable using DL imaging approaches (Broadhead et al., 2016, Pageon et al., 2016, Zhu et al., 2023). Indeed, we have recently used peptide-protein interactions to show the presence of nanoclusters of post-synaptic proteins in fixed synaptosomes (De Moliner et al., 2023). We therefore sought to determine whether such clustering behavior could be observed by H2B in the nucleus of live cells using mLIVE-PAINT.

First, we imaged H2B-101B with LIVE-PAINT, achieving a resolution of <60 nm within 20s of imaging (Figure 2B, Brink, 2022). This allowed us to observe individual H2B proteins organized into distinct densities, reminiscent of nucleosome “clutches” reported previously (Figure 2A, yellow arrows) (Ricci et al., 2015). Subsequent clustering, using DBSCAN (Figure 2C, Pedregosa et al., 2011), highlighted >1000 concentrated patches of localizations covering a mean area of 0.01 m^2^ (Figure 2D) and eccentricity of ∼0.7 (Figure 2F), suggesting that H2B is not evenly distributed in the nucleus (Figure 2B). These results corroborate several previous studies in different cell lines (Wombacher et al., 2010, Lukinavičius et al., 2013, Hauke et al., 2017, Maity et al., 2023). To visualize the distribution of individual H2B molecules, the measured locations were plotted and color-coded based on their local density (Figure 2E, MacGillavry et al., 2013). This process involved calculating the mean density around each molecule by determining the number of neighboring molecules within a radius that was scaled according to the mean density of its cluster. As a result of this analysis, maps displaying the local molecular density within individual clusters were generated. These maps revealed a highly nonuniform distribution of H2B molecules within each cluster, indicating regions of varying density and suggesting a complex organization pattern.

**Figure 2.**
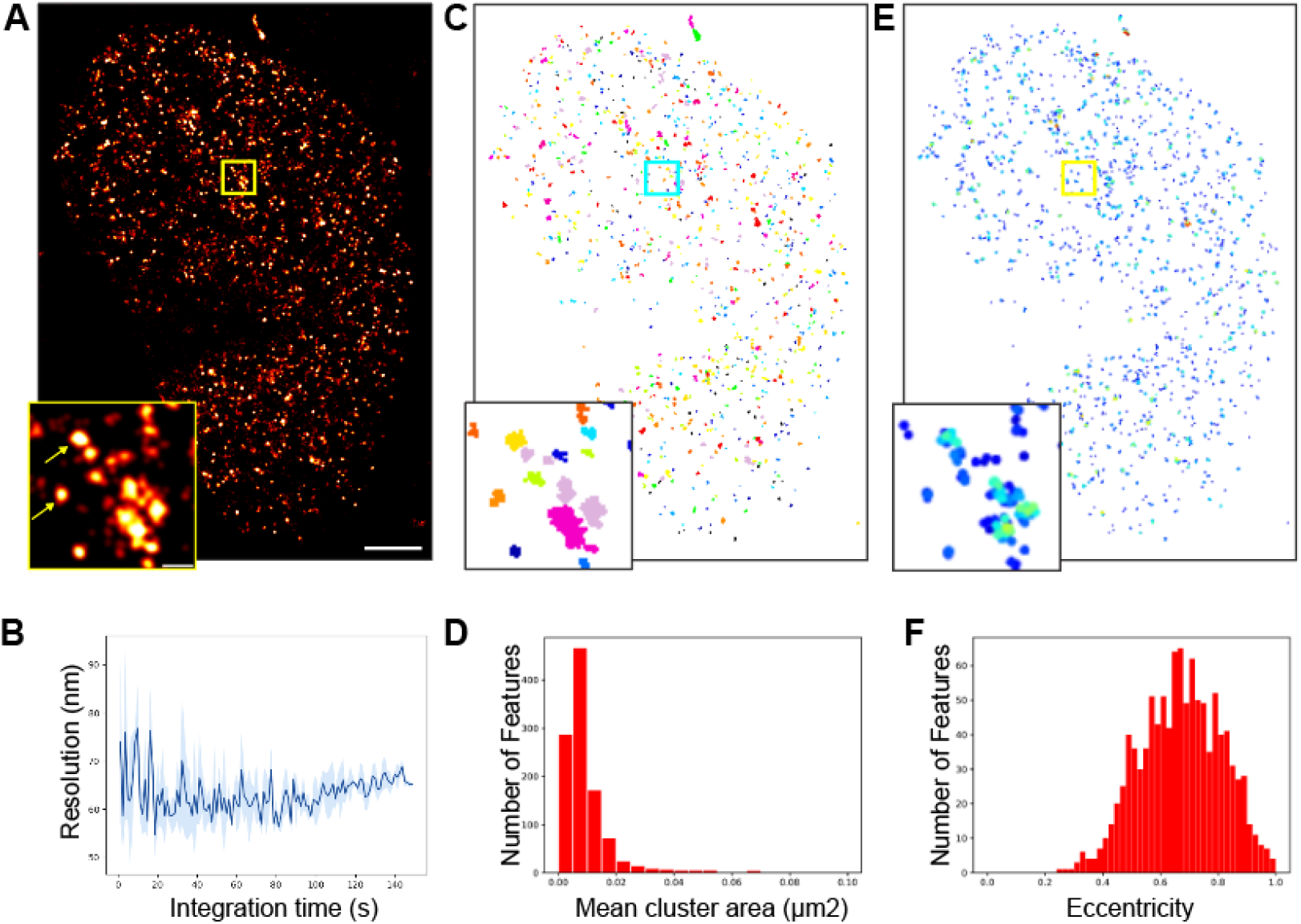
**(a)** SR image of H2B protein located at the nucleus. The imaging was performed over 100 s (2000 frames, 50 ms exposure time) within SH-SY5Y cell stably expressing mNG-101A transfected with H2B-101B. Zoomed in ROI showing distinct clustered localizations. Scale bar for full sized image is 2 m, and 1 m for the zoom-in. **(b)** Resolution response to increasing integration times. Frame subsets were sampled randomly at each integration time for resolution calculation with Fourier Ring Correlation (FRC). Standard deviation error is shown as shaded region (n=3). Precision threshold <30 nm. **(c)** DBSCAN analysis reveals clustered organization of H2B. Rainbow colors distinguish neighboring clusters. **(d)** Histogram of mean cluster areas show areas of 0.01 m^2^ (n=1). **(e)** Nearest neighbor (NN) analysis shows a high local density of proteins within the clusters. **(f)** Distribution of H2B cluster eccentricity (calculated as ratio of major to minor axis lengths, n=1).

### The dynamics of mitochondria can be captured using mLIVE-PAINT

Mitochondria have been imaged extensively using fluorescence microscopy techniques in live mammalian cells (Wurm et al., 2011, Jans et al., 2013, Palmer et al., 2021). Dynamic imaging approaches have also been applied to track fission and fusion events in response to mitochondrial stressors (Guo et al., 2018). In this study, we aimed to show the versatility of mLIVE-PAINT by imaging a dynamic target in the cytoplasm.

Imaging for a period of ∼100 s using TIRF microscopy revealed mitochondria that were motile and underwent changes in position and shape within this timescale (Figure 3A, and SI Movie 1). Subsetting the imaging period into 25 s time-frames and reconstructing the localizations from this period revealed the dynamic structures within the ROI. Comparing this region with the full FOV showed that not all regions were dynamic to the same extent. To better visualize these shape changes over time, we plotted color-coded localizations based on time of acquisition (Figure 3B), which highlighted the spatiotemporal dynamics of mitochondrial networks, especially within the ROI. Higher localisation rates could enable detection of faster dynamics such as fission and fusion events with this technique.

**Figure 3.**
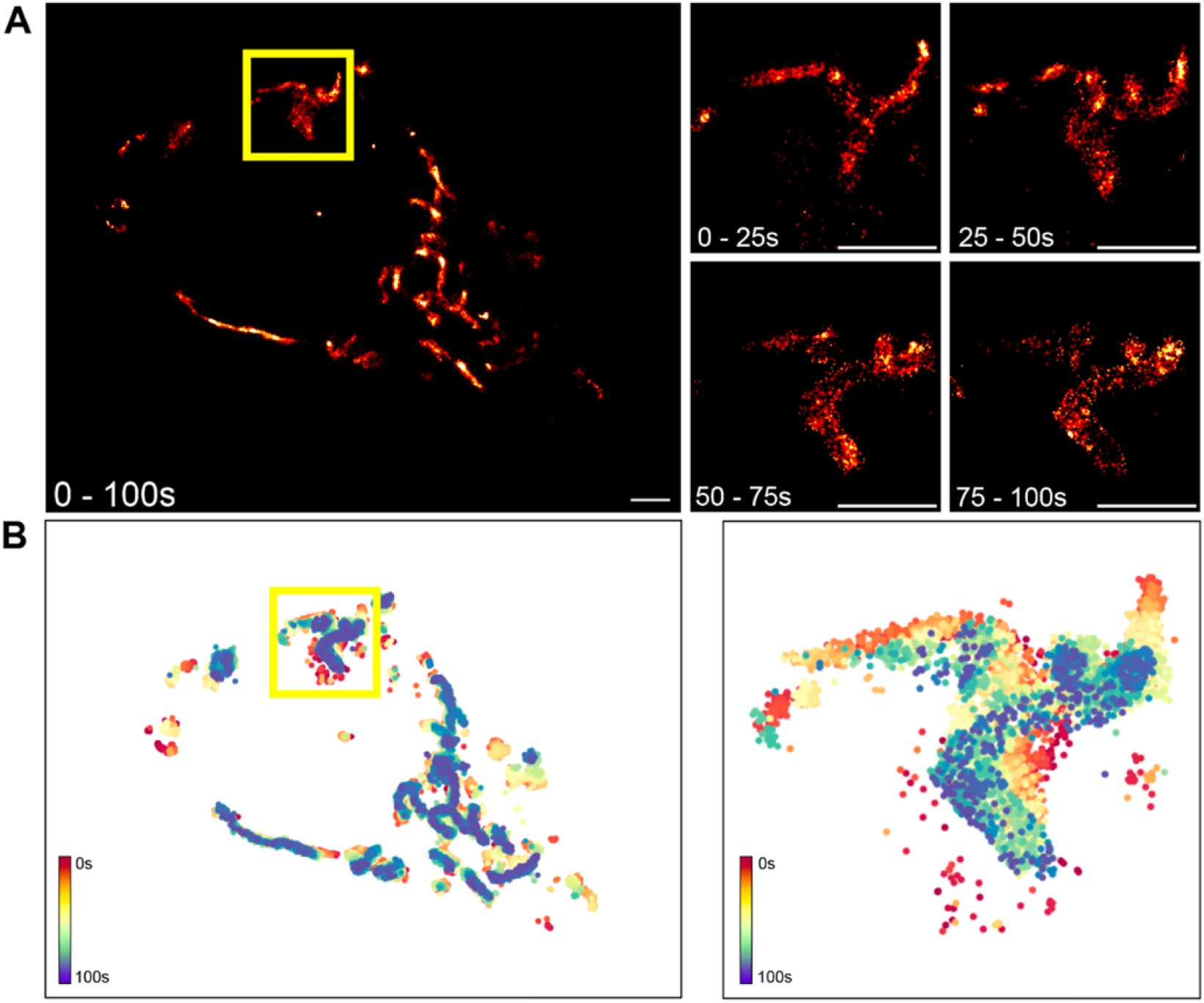
**(a)** SR images of TOM20 imaged over a period of 100 s. 2 μm scale bar. Subsequent panels show a ROI reconstructed with images from discrete time periods within the total 100 s imaging duration. 1 μm scale bar. **(b)** Localisations from the whole FOV and the ROI over the 100 s imaging period, color-coded based on time of acquisition between 0-100 s.

## Discussion

In this study, we have demonstrated that peptide binding pairs can be used for PAINT imaging of proteins in live mammalian cells at the nanometer length scale. Although achieved using the CC peptide pair 101A/B, other peptide pairs are available and would be amenable to this approach, for example the SYNZIP pairs (Thompson et al., 2012) and the tetratricopeptide repeat (TPR) proteins (Jackrel et al., 2010). Indeed, the use of multiple orthogonal pairs would enable multi-target imaging, as we have demonstrated previously in yeast (Gidden et al., 2023). Although other approaches are available for SR imaging in live cells, they each have their own drawbacks. Stimulated emission depletion microscopy (STED, Hell and Wichmann, 1994) requires the use of high laser powers, which can lead to phototoxicity (Vicidomini et al., 2018). Structured illumination microscopy (SIM) uses lower laser excitation powers, but requires the use of expensive equipment and can only achieve resolutions of ∼100 nm (Gustafsson, 2005). Furthermore, photoactivatable localisation microscopy (PALM), STED and SIM all require the direct fusion of the POI to a large fluorescent protein, which can perturb its function and/or localisation (Lisenbee et al., 2003, Agbulut et al., 2006, Hinrichsen et al., 2017, Paine et al., 2021).

In contrast, mLIVE-PAINT is minimally perturbative, and can be used for imaging multiple protein targets at high resolution with the possibility for signal replenishment and relatively low laser powers. By imaging the nuclear protein H2B, we have shown that LIVE-PAINT is a sensitive technique to probe protein location at the nanometer length scale. We also demonstrated that mLIVE-PAINT can measure the dynamics of organelles at the nanometre length scale by imaging TOM20 in live cells.

## Conclusion

In this study, we demonstrate that LIVE-PAINT using the 101A/101B peptide pair is a versatile method to probe dynamic and relatively static protein targets inside live mammalian cells in SR. Compared to existing live-cell SR approaches, LIVE-PAINT presents a minimally perturbative system that preserves protein function and localisation while also minimising phototoxicity. We also showed examples of downstream analyses that can be performed with this data and aim to apply this technique to image difficult-to-tag proteins involved in human disease in the future.

## Materials and methods

### Stable mammalian cell line generation and characterization

All initial plasmid constructs (mNG-101A and POI-101B) were designed with the Rosser Lab using the EMMA cloning system in mammalian expression vectors (Martella et al., 2017). Further cloning and lentivirus production were performed by the Francis Crick Institute’s vectorcore facility using standard protocols. Lentivirus expressing mNeonGreen-101A was added at ∼5 MOI to SH-SY5Y cells grown in T75 flasks at ∼60% confluency. Expression was carried out for 48 h before applying selection at 1.0 mg/ mL geneticin. Media was refreshed every 2-3 days for 11 days. Surviving cells were single cell sorted based on size and positive fluorescence upon doxycycline induction (2 *μ*g/ mL) into 96-well plates. Plates were inspected for surviving single colonies 2 weeks post sort. Cells were then moved to 48-well plates once confluent and treated with doxycycline at 2 *μ*g/ mL to test for expression and clone quality under a confocal microscope. Clones expressing mNG-101A were expanded further and characterized for suitable expression levels by fluorescence under a range of doxycycline concentrations and frozen down for long-term storage.

### Mammalian cell culture and transfection

SH-SY5Y neuroblastoma cell line was cultured using standard ATCC protocols. Briefly, DMEM was supplemented with 10% FBS and 1% Pen/Strep for cell culture (Gibco™, 31966-021). Cells were subcultured ∼ once every three days by detaching with 0.5 mM EDTA and incubating for 5 mins at 37°C before resuspending in DMEM (Gibco, Cat no. 21063-029) and centrifuging at 200 G for 4 mins. Cell pellet was resuspended in fresh media and transferred to a fresh flask (1:6) or counted with Trypan Blue at a 1:1 cell:dye ratio (1:2 dilution) and seeded at 10-12k cells/ well into Ibidi glass-bottom 18-well imaging plates (#81817). Plates were then incubated at 37°C, 5% CO2 for 24 hr.

All transient transfections of POI-101B were carried out using Lipofectamine™ 3000 reagent according to manufacturer’s protocol and as described in more detail in our previous study (Invitrogen™, L3000015). Briefly, 24 h after cell seeding, transfection reagents were made up in Opti-MEM™ (Gibco™, 31985062) and mixed with plasmid DNA of POI-101B at an equivalent of 25 ng DNA/ well and added to cells. Doxycycline was added 24 h after transfection at a final concentration of 200 ng/ mL and plates were imaged 24 hr after that.

### Live-cell confocal microscopy

Cells were plated on uncoated glass ibidi 18-well plates (#81817) and transfected with POI-101B once at 60-70% confluency. 24h post transfection, doxycycline was added at a final concentration of 200 ng/ mL and incubated for 24h before imaging. If staining with live cell mitochondrial and nuclear markers, MitoLite™ FX600 (AATBIO. 22677) and Hoechst 33342 (Thermo Scientific™, 62249) were diluted in DMEM without phenol red (Gibco™, 31053028) and added to cells 30 mins before imaging to achieve final dilutions of 500X and 1000X respectively. Cells were imaged on a commercial spinning-disk confocal system using a 20X or a 40X water-immersion objective.

### TIRF microscopy and super-resolution analysis

Cells were prepared for imaging as described above for confocal microscopy. All TIRF imaging was performed on the ONI Nanoimager (Oxford Nanoimaging Ltd.) equipped with a 100×/1.4 numerical aperture oil immersion objective lens and ORCA-Flash 4.0 V3 scientific complementary metal-oxide semiconductor (CMOS) camera. TIRF angles between 51-53 degrees were used to maximize signal to background. Stage temperature was set to 30°C. The 488 laser line was used to excite the sample between 0.5-2.5 mW. Images were collected continuously at an exposure of 50 ms for a maximum of 4000 frames (200s).

SR image reconstruction was performed as described in our previous study using the FIJI plugin, ThunderSTORM (version dev-2016-09-10-b1, Ovesný et al., 2014), and FRC resolution calculation was performed using the RustFRC python package (Brink, 2022). DBSCAN clustering was performed using the DBSCAN python package (Pedregosa et al., 2011). DBSCAN parameters were set (eps_threshold=0.3, minimum_localisations=5) using the script and results visualized using the matplotlib and seaborn packages on Python. Mitochondria dynamics was visualized using the seaborn package on Python. All scripts used in this study are available at doi: 10.5281/zenodo.11502743.

## Author contributions

Conceptualization: L.R. and M.H.H. Methodology: H.B., Z.G., D.J.K., S.J.R., S.G., L.R. and M.H.H. Investigation: H.B., Z.G., G.V., S.G., L.R., and M.H.H. Supervision: S.G., L.R. and M.H.H. Writing—original draft: H.B. Writing—review and editing: all authors.

## Acknowledgements

The authors would like to thank Molly Strom and Ana Cunha from the Francis Crick Institute’s Viral Vector services for their support in building plasmids and lentivirus used in this study. We would also like to thank the Institute of Regeneration and Repair (IRR) Flow Facility (University of Edinburgh) for their cell sorting expertise used for stable cell line generation and Dr. Justyna Cholewa-Waclaw for her support in the spinning disk confocal imaging experiments. Finally, we thank Drs. Marcus Wilson & Nick Gilbert (University of Edinburgh) for their valuable input on histone protein organization and nuclear structure. The ONI was funded by UCB Biopharma. We also wish to thank Dr. Jim Love for a kind donation to M.H.H.

